# Senescent-like microglia limit remyelination through the senescence associated secretory phenotype

**DOI:** 10.1101/2024.05.23.595605

**Authors:** Phillip S. Gross, Violeta Duran Laforet, Zeeba Manavi, Sameera Zia, Sung Hyun Lee, Nataliia Shults, Sean Selva, Enrique Alvarez, Jason R. Plemel, Dorothy P. Schafer, Jeffrey K. Huang

## Abstract

The capacity to regenerate myelin in the central nervous system (CNS) diminishes with age. This decline is particularly evident in multiple sclerosis (MS), which has been suggested to exhibit features of accelerated biological aging. Whether cellular senescence, a hallmark of aging, contributes to remyelination impairment remains unknown. Here, we show that senescent cells (SCs) accumulate within demyelinated lesions after injury, and their elimination enhances remyelination in young mice but not in aged mice. In young mice, we observed the upregulation of senescence-associated transcripts primarily in microglia after demyelination, followed by their reduction during remyelination. However, in aged mice, senescence-associated factors persisted within lesions, correlating with inefficient remyelination. We found that SC elimination enhanced remyelination in young mice but was ineffective in aged mice. Proteomic analysis of senescence-associated secretory phenotype (SASP) revealed elevated levels of CCL11/Eotaxin-1 in lesions, which was found to inhibit efficient oligodendrocyte maturation. These results suggest therapeutic targeting of SASP components, such as CCL11, may improve remyelination in aging and MS.

## INTRODUCTION

Multiple Sclerosis (MS) is a chronic, autoimmune demyelinating disease, in which immune cells infiltrate the central nervous system (CNS) and destroy myelin and oligodendrocytes, resulting in denuded axons and progressive neurodegeneration^1^. Early in the disease, oligodendrocyte precursor cells (OPCs) migrate to the site of the demyelinated lesions and differentiate into mature myelinating oligodendrocytes, a regenerative process termed remyelination^2^. However, with age, remyelination becomes inefficient^3,4^. This is mirrored in the clinical disease course, where for reasons that remain poorly understood, patients typically transition from Relapse Remitting MS (RRMS) to Secondary Progressive MS (SPMS), characterized by the progressive worsening of symptoms^5–7^. One potential reason for disease progression is the failure of the remyelination process to keep pace with demyelination, leading to progressive axonal dystrophy and neurodegeneration. Notably, therapeutics for MS predominantly rely on modifying the immune system during RRMS, with a limited focus on improving remyelination. Further, most therapeutics become ineffective once the patient progresses to SPMS, which usually occurs during middle age regardless of disease duration^8–10^. Since SPMS onset is associated with increasing age, we hypothesize that the accumulation of senescent cells (SCs) with age limits remyelination, resulting in increased neurodegeneration and clinical deterioration.

SCs are cells locked in permanent cell cycle arrest that become resistant to apoptosis, and release a heterogenous milieu of inflammatory factors termed the senescence associated secretory phenotype (SASP)^11,12^. Often associated with increasing age, these cells have been implicated in multiple age-related neurodegenerative diseases^13,14^. The selective clearance of SCs results in alleviation of disease symptoms and progression^15–17^, extension of lifespan^18,19^, and improved healthspan^20,21^. Interestingly, MS has been shown to be a disease of accelerated aging^22^, and recent evidence of SCs in MS lesions has been reported^23–25^. However, the impact of SCs on remyelination efficiency remains unknown.

Here, we show the upregulation of the senescence marker p16^ink4a^ in microglial cells in demyelinated lesions following acute demyelination and its gradual decline in expression during efficient remyelination in mice. Furthermore, we do not detect obvious cellular senescence in oligodendrocyte lineage cells. The extent of p16+ microglia in demyelinated lesions increases and persists in lesions with advancing age. Using Multiplexed Error-Robust Fluorescence in situ Hybridization (MERFISH) and protein multiplexing, we found that young and aged mouse lesions display transcripts associated with cellular senescence and neuroinflammation, with higher expression in aged mice. Notably, reduction of SCs using genetic and pharmacological methods leads to enhanced remyelination in young and middle-aged mice, but fails to improve remyelination in aged mice. RNA-sequencing and protein multiplexing analyses revealed SC depletion reduced the levels of SASP proteins within aged lesions. We identified CCL11/Eotaxin-1 as one of the major SASP factors affected through SC depletion. Moreover, elevated CCL11 levels correlate with aging and disease severity in MS, and inhibiting CCL11 in mice partially mimics the beneficial effects of SC depletion on remyelination. Our results suggest SCs limit remyelination through the accumulation of SASP in demyelinated lesions, and that inhibition of SASP, such as CCL11, may increase remyelination efficiency by fostering a more favorable regenerative environment.

## RESULTS

### Senescent cells are observed within lesions following demyelination

Senescent cells (SC) have been observed in the brains of individuals with multiple sclerosis (MS), but their impact on the remyelination process remains unclear. Chronic active lesions in MS exhibit increased microglial activation, which is associated with impaired remyelination^24^. Further, it has been shown that p16, a marker of cellular senescence, could be observed in microglia within the brains of individuals with MS^24^. Re-analysis of a previously published single nucleus RNA sequencing (snRNA-seq) database of MS lesions^34^ revealed an elevation of *Cdkn2a*, the gene encoding p16, in both chronic active and inactive MS lesions compared to control brain tissues (**Extended Data Fig. 1a,b**). To determine if SCs are detectable in demyelinated lesions in mice, focal demyelination, by injection of lysophosphatidylcholine (LPC), was performed on spinal cord white matter of young adult mice (3 months old). This approach induces acute demyelination and subsequent remyelination in a predictable manner (**Fig. 1a**). Staining of lesioned spinal cord tissues for senescence associated ß-galactosidase (B-gal) activity revealed a significant increase in B-gal+ cells at 5 days post lesion (dpl) compared to non-lesioned control white matter tissue (**Fig. 1b,c**). Moreover, RT-qPCR analysis of demyelinated lesions and non-lesioned control at 5 dpl revealed demyelinated lesions exhibited upregulation of *p16/Cdkn2a* and *Rb1,* associated with cellular senescence (**Fig. 1d,e**), downregulation *Tert*, associated with a youthful cell state (**Fig. 1f**), and upregulation of *Apoe* and *Trem2,* associated with disease-associated microglia (DAM) (**Fig. 1g,h**). Additionally, there was a downregulation of *p21/Cdkn1a* and no difference in *Trp53* and *E2f1* (**Extended Data Fig. 1c-e**). Furthermore, analysis of mRNA transcripts from bulk RNA-sequencing of dissected lesions and non-lesioned tissues at 20 dpl revealed that 23 out of 103 genes from the SenMayo gene panel, a dataset of differentially expressed genes in senescent cells^35^, were significantly upregulated in lesions compared to non-lesioned tissues (**Fig. 1i**).

**Fig. 1.**
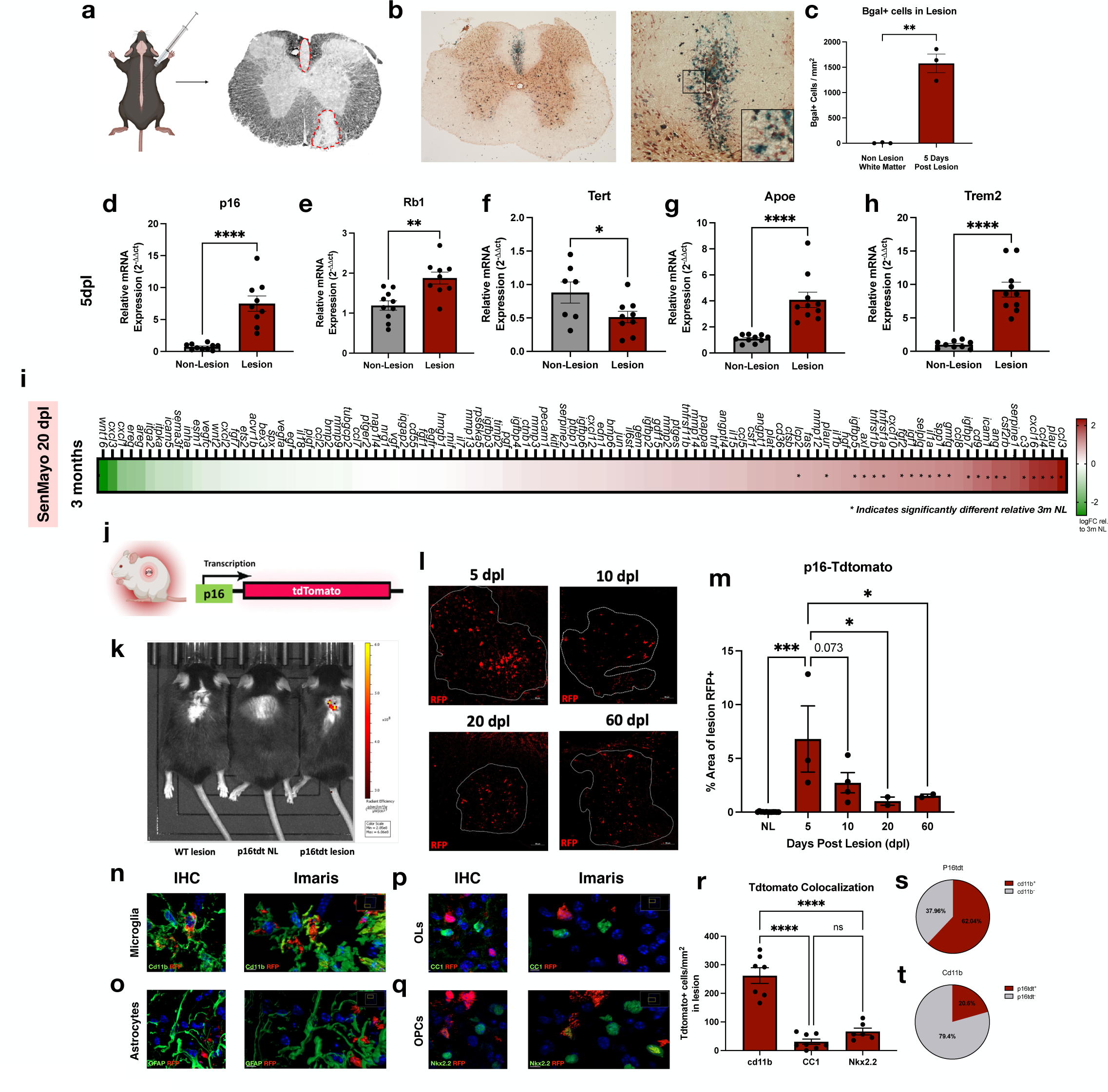
Senescent cells are observed within lesions following demyelination. **a,** Schematic of lysophosphatidylcholine (LPC) induced demyelination. **b,** Enzymatic staining of senescence associated ß-galactosidase of a dorsal demyelinated lesion at 5 dpl in young adult mouse. **c,** Quantification of ß-gal+ cells in b (n= 3). **d-h,** RT-qPCR for selected senescence genes from dissected lesion (n= 9-10). **i,** Heatmap depicting SenMayo gene expression in young lesions relative to young non-lesions (n=3). Color represents LogFC relative to young non-lesion and * indicates significantly differentially expressed. **j,** Schematic of p16-tdTomato mouse. **k,** In Vivo Imaging System (IVIS) bioluminescence of controls and p16tdt lesions (n= 1-2 per group). **l,** Immunofluorescent staining of p16tdt+ cells throughout remyelination. **m,** Quantification of k (n= 2-4 per timepoint). **n-q,** Immunofluorescent staining and Imaris 3D reconstruction of p16tdt and Cd11b, GFAP, CC1, and Nkx2.2 colocalization. **r,** Quantification of m-p (n= 6-8). **s,t**, Percentages of p16tdt+ cells Cd11b+ and Cd11b+ cells p16tdt+. Data are mean ± SEM (**c-h,m,r**). Gene expression levels were expressed using the 2^−ΔΔCt^ method normalized to non-lesion tissue (**d-h**). *P* values derived from two-tailed unpaired Student’s t-tests (**c-h**), one-way ANOVA with Tukey corrected multiple comparisons (**m,r**), or the Wald test (**i**).

To quantify the level of SCs in lesions during remyelination, we performed LPC-induced demyelination on p16-tdTomato (p16tdt) mice, which express a tdTomato reporter under the endogenous promotor of p16^ink4a^, a key gene in senescence programming^26^ (**Fig. 1j**). We first determined if the p16tdt reporter is detectable in lesions after LPC-induced demyelination in young mice. Using IVIS imaging, we observed increased bioluminescence in spinal cord lesions of p16tdt mice compared to LPC-lesioned wildtype or sham lesioned p16tdt mice (**Fig. 1k** and **Extended Data Fig. 1f,g**). We next examined p16tdt reporter expression at 5 dpl (corresponding with inflammation and OPC recruitment), 10 dpl (corresponding with inflammation resolution and OPC differentiation), 20 dpl (corresponding with completion of remyelination), and 60 dpl (corresponding with extended oligodendrocyte myelination)^27,36^ (**Fig. 1l**). We found that compared to non-lesioned control white matter, which exhibit very low p16tdt expression (**Extended Data Fig. 1h**), demyelinated lesions showed a substantial increase in p16tdt expression at 5 dpl followed by a gradual decrease at 10, 20, and 60 dpl (**Fig. 1m**). The pattern of p16tdt distribution in lesions over the course of remyelination was further verified by immunostaining analysis of lesions for p16 and p21 expression (**Extended Data Fig. 1i-k**).

To determine which cell population(s) exhibit cellular senescence in demyelinated lesions, we co-stained the p16tdt lesions with markers of myeloid cells (CD11b), astrocytes (GFAP), mature oligodendrocytes (CC1) and OPCs (Nkx2.2) (**Fig. 1n-q**). We found p16tdt expression predominantly co-localized with CD11b+ cells in lesions (**Fig. 1r** and **Extended Data Fig. 2a**) accounting for 62% of the p16tdt+ cells (**Fig. 1s**), and 20% of the myeloid cells in the lesion (**Fig. 1t**). Moreover, co-immunostaining analysis for p16tdt and SALL1, a microglia-associated marker^37^, suggest the senescent myeloid cells are likely derived from microglia (**Extended Data Fig. 2b,c**). Furthermore, we observed minimal expression of p16tdt in oligodendrocyte progenitors and oligodendrocytes, indicating oligodendrocyte lineage cells do not undergo cellular senescence within demyelinated lesions (**Fig. 1r** and **Extended Data Fig. 2a**). Taken together, our findings suggest that cellular senescence increases primarily in microglia within demyelinated lesions, which naturally decreases during efficient remyelination in young adult mice.

### Persistence of senescent cells in lesions are associated with inefficient remyelination in aged mice

Since remyelination becomes less efficient with age, we next asked if the presence of SCs in demyelinated lesions increases with age and potentially contributes to remyelination impairment. To assess the level of cellular senescence in lesions of aged mice compared to young mice, we performed spatial gene expression profiling of lesioned spinal cord tissues in 3 months (young) and 18 months (aged) old mice at 5 days post lesion (dpl) using Multiplexed Error-Robust Fluorescence In Situ Hybridization (MERFISH) with a customized panel designed to characterize senescence associated gene expression (**Fig. 2a**). Focal lesion was delineated by the hypernuclear DAPI staining in both the young and aged mouse spinal cord sections (**Fig. 2b** and **Extended Data Fig. 3a**). Analysis of distinct cell populations within the outlined lesioned area through UMAP clustering of cell-type associated gene markers revealed all major cell types found in the spinal cord, including microglia, macrophages, OPCs, oligodendrocytes, astrocytes, and endothelial cells (**Fig. 2c,d** and **Extended Data Fig. 3b-d**). The relative percentage of cells identified in lesions differed substantially from non-lesions. In lesions, an expansion of macrophages and microglia, along with a reduction in oligodendrocytes, observed at both ages (**Extended Data Fig. 3e**). Moreover, demyelinated lesions exhibited a significant increase in transcripts associated with senescence, such as *Lgals3*^38,39^, *Spp1*, and *Ctsb*, compared to non-lesioned tissues (**Extended Data Fig. 3f**). We also observed the downregulation of *Cdkn1a* (p21) in lesions, consistent with our observations in Extended Fig. 1. When comparing lesions between aged and young mice, we found that lesions from aged mice exhibited an upregulation of a number of genes associated with cellular senescence and microglial activation, including *Spp1* and *C3*, both part of the SenMayo and DAM genesets, *Irf7*, which has been shown to induce senescence^40^, *CCnd1*, which is hyper-expressed in SCs^41^, and *Efnb3*, which is necessary for SC survival^42^ (**Fig. 2e,f**).

**Fig. 2.**
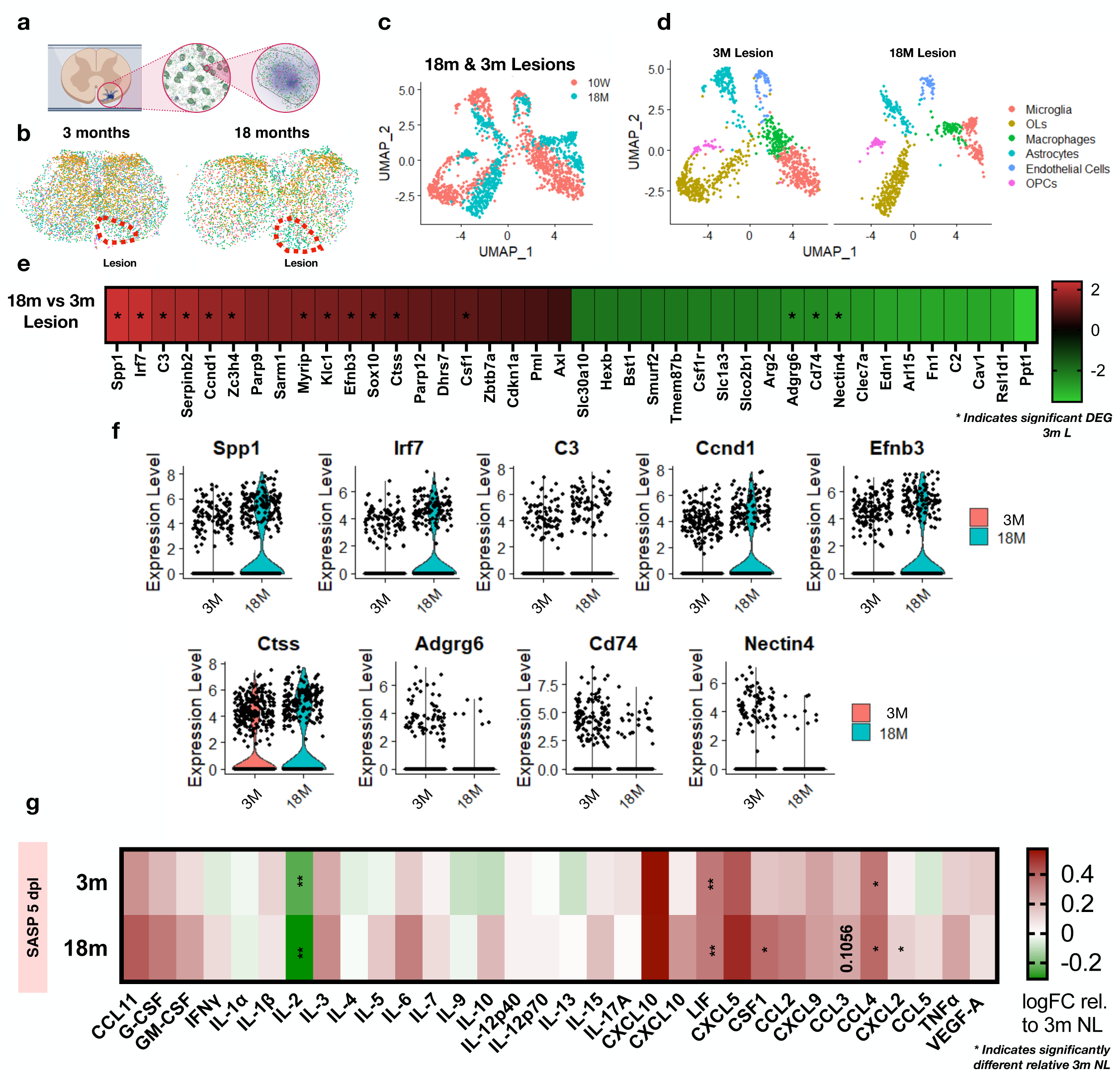
Increased senescent markers and SASP proteins characterize aged lesion after demyelination. **a,** Schematic of MERFISH spatial transcriptomics. **b,** Demyelinated lesions were traced in young (3 months) and aged (18 months) mice at 5 dpl. **c,** Uniform manifold approximation and projection (UMAP) visualization of pooled lesions. **d,** UMAP visualization of lesion cell types in young and aged lesions (n=2 per group). **e,** Heatmap depicting top differentially expressed genes between young and aged lesions at 5 dpl. Color represents LogFC relative to young lesion and * indicates significantly differentially expressed. **f,** Expression levels of selected genes from e. **g,** Heatmap depicting SASP protein levels in young and aged lesions relative to young non-lesioned white matter at 5 dpl (n=3 per group). Color represents LogFC relative to young non-lesion and * indicates significantly differentially expressed. Data are mean ± SEM (**f**). *P* values derived from two-way ANOVA with FDR corrected multiple comparisons (**g**) or the Wald test (**e,f**).

To examine the presence of senescence associated secretory phenotype (SASP) factors in lesions, we performed multiplex protein analysis of dissected lesions and non-lesioned tissues at 5 dpl in young and aged mice. In both age groups, we found that lesioned tissues exhibited a decrease in IL-2 (**Fig. 2g**), an increase in LIF, and an increase in CCL4 compared to non-lesioned control tissues. Moreover, in aged lesions, an increase in CSF1, CXCL2, and IL-5 was observed (**Extended Data Fig. 3g-i**). Taken together, these data indicate that at 5 dpl, aged mice exhibit increased cellular senescence accompanied by an elevation in SASP factors in demyelinated lesions compared to young mice.

To determine if the presence of SCs in lesions correlates with inefficient remyelination in aged mice, we compared the expression of p21 in the lesions of young and aged mice. We observed that aged mice exhibited an increased level of p21 expression compared to young mice at 20 dpl (**Fig. 3a,b**), which persisted in lesions at 60 dpl (**Fig. 3c,d**). The sustained elevation of p21 coincided with a reduction in OPC differentiation and mature oligodendrocytes within the lesion at 20 dpl (**Fig. 3e-h** and **Extended Data Fig. 4a,b**). To further characterize the difference between young and aged mouse lesions, we dissected the lesions from both young and aged mice for bulk-RNA sequencing at 20 dpl (**Fig 3i**). We identified 434 differentially expressed genes between the lesions of young and aged mice (**Fig. 3j**). Using the SenMayo gene panel, we found that 14 senescence associated transcripts were significantly differentially expressed in lesions of aged mice compared to young mice (**Fig. 3k**). Moreover, we found that transcripts corresponding to damage associated microglia (DAM) were also elevated in aged mice compared to young mice (**Extended Data Fig. 4c**). Immunostaining analysis of IBA1 and CD68 for activated microglia confirmed aged mice exhibited sustained microglial activation in lesions compared to young mice at 20 dpl, but their expressions levels were found to decrease to similar levels as young mice by 60 dpl (**Extended Data Fig. 4d-g**). We also examined the level of SASPs in the lesions at 20 dpl, and found that aged mice exhibited increases in CSF1, CCL4, CCL5, CCL11, CXCL1, CXCL2, and a decrease in IL2 compared young mice (**Fig. 3l**). These findings indicate that demyelinated lesions of aged mice exhibit prolonged SC and DAM activity, corresponding with the failure of OPCs to efficiently differentiate into remyelinating oligodendrocytes.

**Fig. 3.**
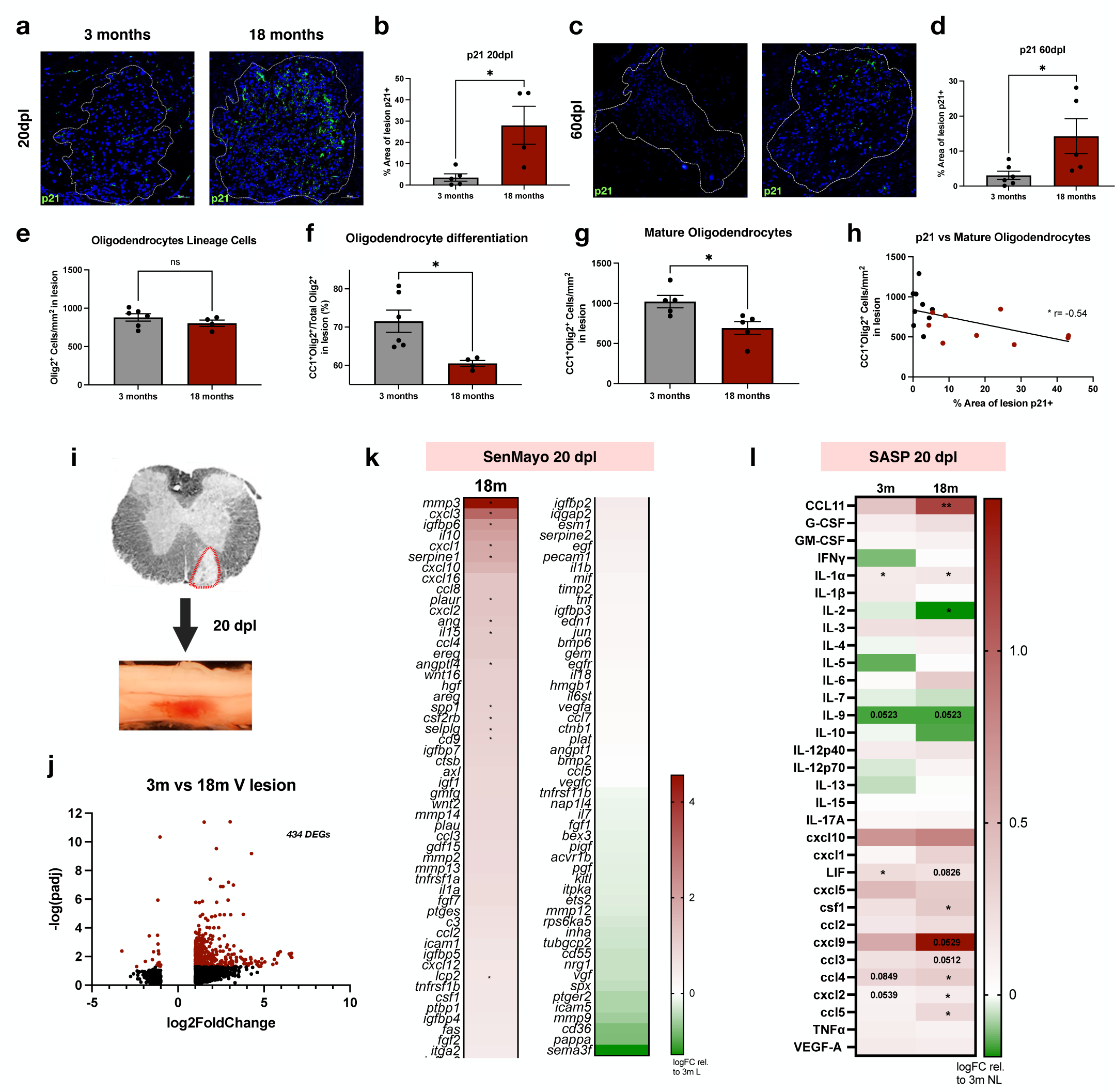
Persistence of senescent cells in lesions are associated with inefficient remyelination in aged mice. **a,** Immunofluorescent staining of p21 in young and aged (18 months) lesions at 20 dpl. **b,** Quantification of p21+ cells in a (n= 4-5 per age). **c,** Immunofluorescent staining of p21 in young and aged lesions at 60 dpl. **d,** Quantification of p21+ cells in c (n= 5-6 per age). **e,** Quantification of *Olig2* in young and aged lesions (n= 4-6 per age). **f,** Quantification of percentage of differentiated oligodendrocytes in young and aged lesions (n= 4-6 per age). **g,** Quantification of mature oligodendrocytes and oligodendrocyte lineage cells with *CC1* and *Olig2* in young and aged lesions (n= 5 per age). **h,** Correlation between p21+ staining and mature oligodendrocytes (*CC1+Olig2+*) in the lesions. **i,** Experimental schematic of lesion microdissection. **j,** Volcano plot of significantly differentially expressed genes between young demyelinated lesions and aged lesions (n=3 per group) at 20 dpl. Genes with adjusted p-values <0.05 and absolute log2 fold changes >1 were called as differentially expressed genes for each comparison. **k,** Heatmap depicting SenMayo gene expression in aged lesions relative to young lesions (n=3 per group). Color represents LogFC relative to young lesion and * indicates significantly differentially expressed **l,** Heatmap depicting SASP protein levels in young and aged lesions relative to young non-lesioned white matter (n=3 per group). Color represents LogFC relative to young non-lesion and * indicates significantly differentially expressed. Data are mean ± SEM (**b,d-g**). *P* values derived from two-tailed unpaired Student’s t-tests (**b,d-g**), simple-linear regression (**h**), two-way ANOVA with FDR corrected multiple comparisons (**l**) or the Wald test (**j,k**).

### Senescent cell elimination increases remyelination in an age-dependent manner

To examine the impact of SC depletion on remyelination efficiency, we performed LPC-induced demyelination on INK-ATTAC (IA) mice at 3 months (young), 12 months (middle-aged) or 18 months (aged) old. These mice transgenically express a FK-binding protein fused to caspase-8 under the p16^ink4a^ promotor, enabling the induction of apoptosis in p16-expressing SCs upon systemic injection of the synthetic drug AP20187 (AP)^18^ (**Fig. 4a**). We administered AP (2 mg/kg) or vehicle (control) twice a week in IA mice until 20 dpl following LPC-induced demyelination before the mice were perfused for analysis (**Fig. 4b**). To examine the effect of AP treatment on remyelination, immunostaining analysis for oligodendrocyte lineage cells was performed at 20 dpl. In young mice, we observed an increase in Olig2+ oligodendrocyte lineage cells and CC1+ mature oligodendrocytes in lesions after AP treatment compared to control (**Fig. 4d-f**). Moreover, scanning electron microscopy (SEM) analysis of lesions revealed AP treatment modestly, but significantly, increased the extent of remyelination compared to vehicle treated control (**Extended Data Fig. 5a-c**). In middle-aged mice, we found that AP treatment did not significantly affect the density of oligodendrocyte lineage cells, but resulted in an increase in mature oligodendrocytes and myelin basic protein (MBP) compared to vehicle treated controls (**Fig. 4g-i** and **Extended Data Fig. 5d,g,h**). We performed similar experiments in a more clinically relevant model by administering the senolytic cocktail of dasatinib and quercetin (D/Q), which are currently being used in several clinical trials to deplete senescent cells^42,43^. We found that two sets of 3 consecutive doses of D/Q (**Fig. 4c**) treatment also resulted in increased MBP expression in middle-aged lesions compared to control with no difference in oligodendrocyte lineage cells, OPC differentiation, or mature oligodendrocytes (**Fig. 4m-o** and **Extended Data Fig. 5f,k,l**). Intriguingly, in aged mice, AP treatment did not affect the density of oligodendrocyte lineage cells, percentage of mature oligodendrocytes, or the level of MBP in lesions compared to vehicle treated controls (**Fig. 4j-l** and **Extended Data Fig. 5e,i,j**). These results suggest that the depletion of SCs increases oligodendrocyte maturation and remyelination in young and middle-aged mice, but is unable to do so in aged mice.

**Fig. 4.**
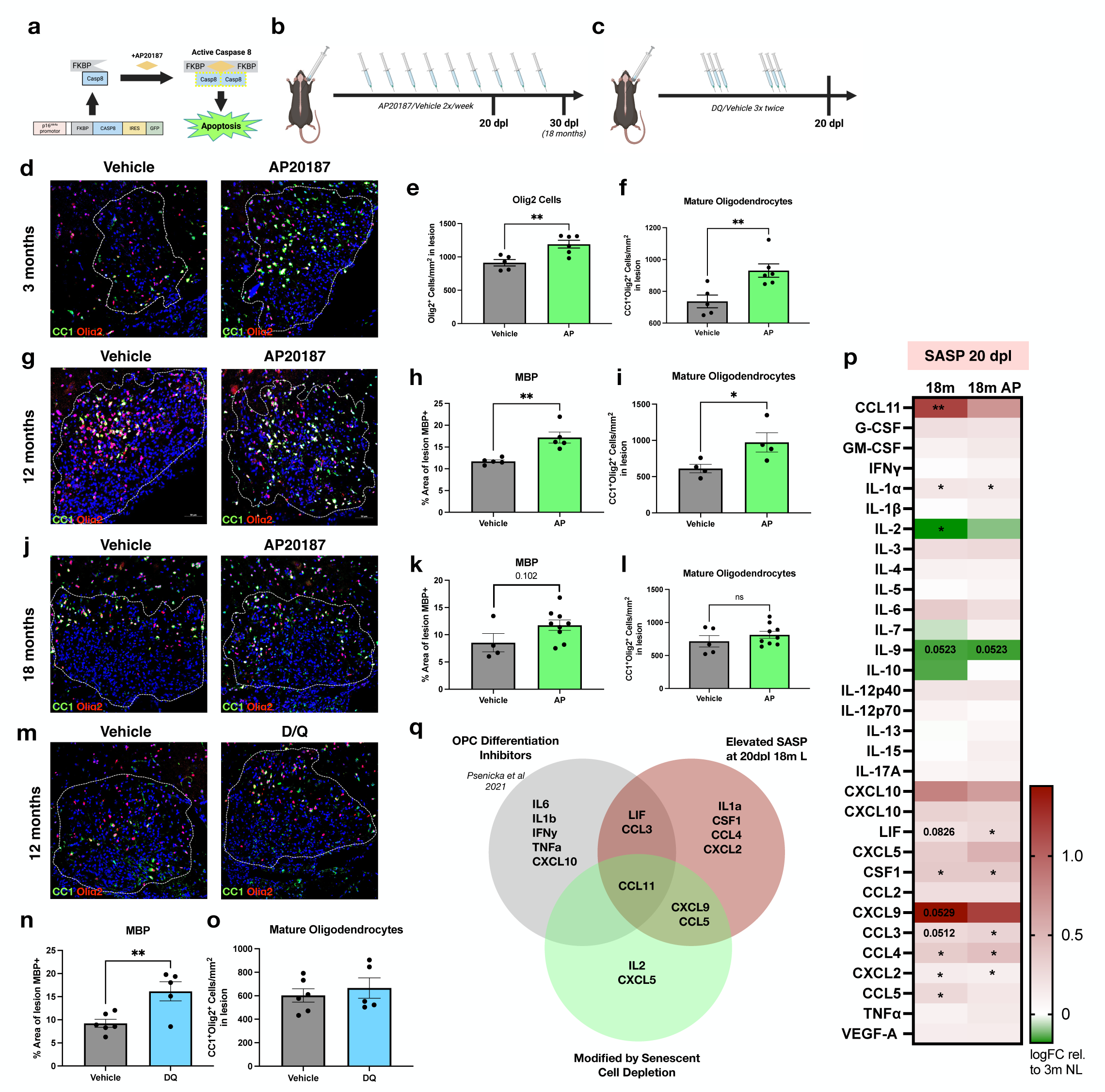
Senescent cell elimination increases remyelination in young but not aged mice. **a,** Schematic of INK-ATTAC mouse model. **b-c,** Experimental schematics of vehicle, AP20187 (2 mg/kg) and Dasatinib (5 mg/kg)/Quercetin (50 mg/kg) treatments. **d,** Immunofluorescent staining of mature oligodendrocytes and oligodendrocyte lineage cells with *CC1* and *Olig2* in young (3 months) vehicle and AP treated lesions at 20 dpl. **e,** quantification of *Olig2+* cells in d (n=5-6 per group). **f,** quantification of *CC1+Olig2+* cells in d (n=5-6 per group). **g,** Immunofluorescent staining of mature oligodendrocytes and oligodendrocyte lineage cells with *CC1* and *Olig2* in middle-aged (12 months) vehicle and AP treated lesions at 20 dpl. **h,** quantification of *MBP+* area in extended data fig. 5d (n=5 per group). **l,** quantification of *CC1+Olig2+* cells in g (n=5 per group). **j,** Immunofluorescent staining of mature oligodendrocytes and oligodendrocyte lineage cells with *CC1* and *Olig2* in aged (18 months) vehicle and AP treated lesions at 30 dpl. **k,** quantification of *MBP+* area in extended data fig. 5e (n=4-9 per group). **l,** quantification of *CC1+Olig2+* cells in j (n=5-9 per group). **m,** Immunofluorescent staining of mature oligodendrocytes and oligodendrocyte lineage cells with *CC1* and *Olig2* in middle-aged vehicle and D/Q treated lesions at 20 dpl. **n,** quantification of *MBP+* area in extended data fig. 5f (n=5-6 per group). **o,** quantification of *CC1+Olig2+* cells in m (n=5-6 per group). **p,** Heatmap depicting SASP protein levels in aged vehicle treated and aged AP treated lesions relative to young non-lesioned white matter (n=3 per group). Color represents LogFC relative to young non-lesion and * indicates significantly differentially expressed. **q,** Venn-diagram of elevated SASP factors in the aged demyelinated lesions, SASP factors changed with clearance of senescent cells, and proteins known to restrict OPC differentiation. Data are mean ± SEM (**e,f,h,i,k,l,n,o**). *P* values derived from two-tailed unpaired Student’s t-tests (**e,f,h,i,k,l,n,o**), or two-way ANOVA with FDR corrected multiple comparisons (**p**).

We next sought to determine the mechanism by which senolytics confer increased remyelination in young and middle-aged mice but not in aged mice, despite the presence of elevated senescence signature in the lesions of aged mice. To compare SC activities in lesions between young and aged mice treated with either vehicle or AP, we conducted bulk RNAseq analysis of lesion tissues at 20 days post-lesion (dpl). Notably, while there were 434 differentially expressed genes between young and vehicle-treated aged lesions (**Fig. 3j**), we observed only 70 differentially expressed genes between young and aged mouse lesions after AP treatment (**Extended Data Fig. 6a**). Furthermore, using the SenMayo gene panel, we observed that while 14 senescence-associated transcripts were significantly differentially expressed in lesions of aged mice compared to young mice, there was no significant differential expression of SenMayo genes in AP-treated aged lesions compared to young lesions (**Extended Data Fig. 6b**). Additionally, no differential expression of DAM genes was observed in AP treated aged lesions compared to young lesions (**Extended Data Fig. 6c**). These results suggest that in contrast to lesions in aged mice, which exhibit sustained upregulation of senescence and DAM associated transcripts at 20 dpl, AP treatment significantly reduces these transcripts to levels similar to those in young mice. We also determined if cellular senescence in lesions is regulated at the protein level by measuring the level of SASP proteins in lesions at 20 dpl in aged mice with or without AP treatment. Aged mice were used due to their prolonged remyelination period and amplified senescence phenotype, where protein differences would be more readily captured. Following AP treatment of aged mice, we observed a decrease in multiple SASP factors in lesions, including CCL11, CXCL9, and CCL5, and an increase in IL-2, which were no longer differentially expressed compared to control non-lesion tissue (**Fig. 4p**). These findings suggest AP treatment reduces SC activity in aged mouse lesions despite the lack of significant improvement in remyelination. Possible explanations for the failure to increase remyelination in aged mice are that some SASP factors may remain in lesions after AP treatment, thereby preventing oligodendrocyte differentiation, or that SC reduction/depletion alone is insufficient to improve remyelination in an already aged CNS environment that has multiple inhibitory factors to regeneration.

### CCL11 is a SASP factor contributing to remyelination impairment

To identify SASP factors that may contribute to remyelination impairment in aged mice, we compared SASP factors elevated in aged lesions with those modified by AP-mediated SC depletion. This narrowed our search to CCL5, CCL11, and CXCL9. When these factors were cross-referenced against proteins known to inhibit OPC differentiation^44^, CCL11 emerged as a potential mechanistic target (**Fig. 4q**). CCL11/Eotaxin1 is a chemokine that selectively recruits eosinophils and is elevated with age^45^. However, recently it was also shown to affect the central nervous system in multiple ways: CCL11 efficiently crosses the blood brain barrier^46^, and has been implicated in neuroinflammatory disorders^47,48^ and impaired neurogenesis^48–50^. Moreover, OPCs have been suggested to express the primary CCL11 receptor, CCR3^51^, and elevated CCL11 levels are associated with MS disease duration and severity^52,53^. These observations, combined with our findings, suggests CCL11 may play a central role in age-associated remyelination delay.

To determine if CCL11 is detected in SCs, we examined the lesions of p16tdt mice following LPC-induced demyelination and observed a strong co-localization of CCL11 with p16tdt (**Fig. 5a**). We also found that CCL11 co-labeled Iba1+ microglia in lesions of both young and aged mice. Moreover, the expression level of CCL11 was significantly higher in lesions of aged mice compared to young mice (**Fig. 5b-c**) and was barely observed in non-lesioned control tissues of both mouse groups (**Extended Data Fig. 7a**). We also analyzed our SASP protein multiplex data of demyelinated lesions at 20 dpl and found that CCL11 was upregulated in the lesions of aged mice compared to those of young mice (**Fig. 5d**). Furthermore, AP treatment reduced its expression in aged mouse lesions (**Fig. 5d**).

**Fig. 5.**
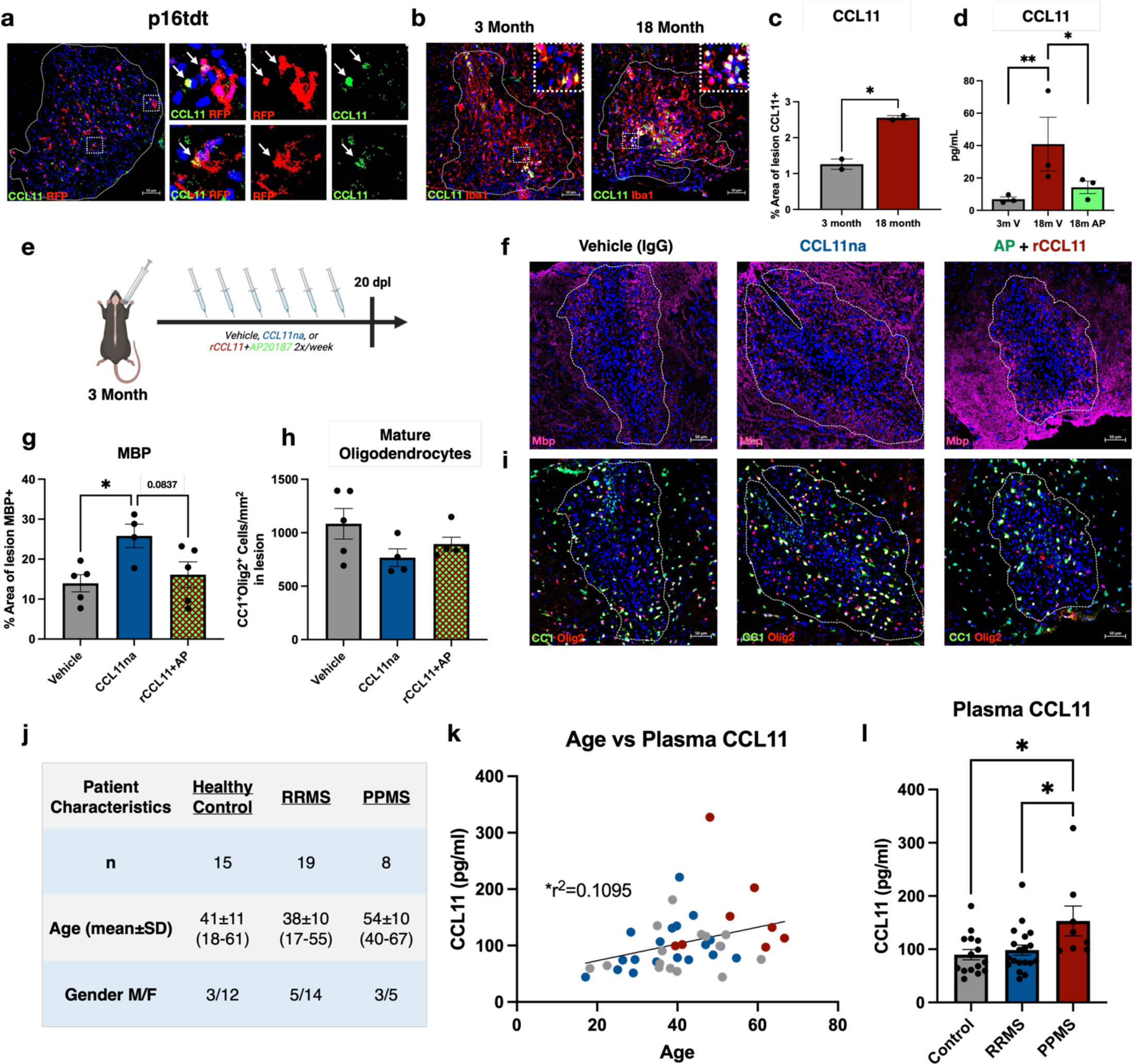
CCL11 is a SASP factor contributing to remyelination impairment. **a,** Immunofluorescent staining of p16tdt+ cells (*RFP*) and *CCL11* in middle-aged (12 months) mouse demyelinated lesions. Arrows indicate p16tdt+CCL11+ colocalization. **b,** Immunofluorescent staining of microglia (*IBA1*) and *CCL11* in young (3 months) and aged (18 months) demyelinated lesions. **c,** Quantification of *CCL11* in b (n= 2 per group). **d,** Protein levels of CCL11 in young, aged vehicle treated, and aged AP treated lesions at 20 dpl (n= 3 per group). **e,** Experimental schematic of IgG vehicle (50 ug/kg), CCL11 neutralizing antibody (50 ug/kg), or AP20187 (2 mg/kg)/recombinant CCL11 (10 ug/kg) treatment of young lesions. **f,g,** Immunofluorescent staining of mature oligodendrocytes with *CC1* and *Olig2* and myelin with *MBP* in vehicle, CCL11na, and rCCL11/AP treated lesions at 20 dpl. **h,i,** Quantification of *MBP*+ area and CC1*+Olig2+* cells in g,h (n=4-5 per group). **j,** Characteristics of human control and MS plasma samples used. **k,** Correlation between age and CCL11 plasma concentration across samples. **l,** CCL11 plasma concentrations in healthy control, Relapse Remitting MS, and Primary Progressive MS samples (n= 8-19 per group, uncorrected for age). Data are mean ± SEM (**c-d,h,i,l**). *P* values derived from two-tailed unpaired Student’s t-tests (**c**), the Wald test (**d**), one-way ANOVA with Tukey corrected multiple comparisons (**h,i,l**), one-way ANOVA with FDR corrected multiple comparisons (**e**), or simple-linear regression (**k**).

To directly test the effects of CCL11 on remyelination *in vivo*, we inhibited CCL11 in young mice by administering a CCL11 neutralizing antibody (CCL11na) systemically twice per week throughout remyelination (**Fig. 5e**). At 20 dpl, we observed an increase in MBP staining in the lesions of the CCL11na treated group compared to the isotype control (**Fig. 5f,g**). However, CCL11na administration did not seem to have an effect on the number of mature oligodendrocytes in lesions (**Fig. 5h,i**). Interestingly, artificially increasing CCL11 levels through the administration of recombinant CCL11 protein (rCCL11) alongside AP treatment in IA mice prevented the positive effects of AP treatment on remyelination (**Fig. 5g,h**). Taken together, our results suggest the SASP factor CCL11 contributes to the inhibitory effects of SCs on remyelination, and that depletion of CCL11 enhances oligodendrocyte maturation by increasing MBP expression.

To validate our findings in a clinical context, we also assessed the concentration of CCL11 in the plasma of individuals with MS and healthy controls (**Fig. 5j**). We found a significant association between age and CCL11 concentration across all samples (**Fig. 5k**). Interestingly, this increase in CCL11 concentration with age also paralleled disease progression (**Fig. 5l**). These results indicate CCL11 level increases with age and disease progression, and suggests inhibition of CCL11 may prevent senescent cell-mediated remyelination failure in MS.

## DISCUSSION

Aging is the number one risk factor for most neurodegenerative disease progression, and with it comes the accumulation of senescent cells. While the role of SCs in these diseases is beginning to be understood, their impact on remyelination in MS is still poorly defined. Here, we report the emergence of senescent-like microglial cells following demyelination, with gradual clearance throughout remyelination. We also find this phenomenon increases with age, which is known to be associated with reduced capacity to remyelinate. Previous studies have shown the accumulation of SCs occurs in injured, or damaged tissue, even in young animals, often as a stress-induced response^54,55^. These SCs have been found to exhibit beneficial effects for regeneration, and become damaging only if unresolved in a timely manner, such as in the context of aging^56^. Other studies have shown contradictory evidence where this SC response limits regeneration throughout repair and regenerative processes and offers no benefits^57,58^. While the beneficial or detrimental effects of damage-induced SCs are likely cell-type and context dependent, we show here that in the context of remyelination, senescent microglia limit remyelination. In an age dependent manner, we observe an increased and prolonged senescence response to demyelination, possibly due to the failure of the aged microglia to clear the SCs as efficiently^59^. Clearance of SCs through genetic and pharmacologic methods resulted in increased numbers of mature oligodendrocytes and improved remyelination in lesions of young and middle-aged mice.

We also showed that multiple SASP factors are dysregulated in aged mouse lesions and were returned to younger levels with senescent cell depletion. In particular, we found that CCL11/Eotaxin-1 is one of the most drastically altered SASP proteins after SC depletion. CCL11 has previously been shown to be associated with aging^45,49^ and MS disease severity^52,53^, to impair neurogenesis and myelination^48–50^, and to cause cognitive dysfunction^47,48^. Following CCL11 inhibition during remyelination in young mice, we observed similar beneficial remyelinating effects as when senescent cells were eliminated through AP treatment. Interestingly, the elevation of plasma CCL11 level was observed in MS patient with age, concurrent with disease progression. Since MS exhibits features of accelerated aging, it is possible that elevated levels of SCs with chronic disease result in increased SASP, including CCL11, which inhibit remyelination and promote the deterioration of symptoms.

However, we found that SC depletion did not significantly improve remyelination in aged mice, despite modifying the lesion environment. This limited effect may be due to incomplete clearance of SCs or lingering SASP production in the aged lesions. Indeed, AP treatment modified SASP profiles within the aged lesions but several factors remained elevated (**Fig. 4p**). Additionally, it is likely that the presence of SCs is only one of multiple factors limiting remyelination in the aged nervous system. Other factors known to change with age include increased chronic inflammation^60^, modified niche stiffness^61^, and inefficient myelin debris removal and processing by microglia and macrophages^62^. Similarly, changes in OPCs with age result in reduced recruitment^63^ and limited intrinsic capacity to differentiate^62,64^, further limiting remyelination.

The cause of senescent microglia accumulation following demyelination has yet to be established. Recent reports suggest substantial overlap in transcriptional profiles between disease associated microglia (DAM), white matter associated microglia (WAM), and senescent cells^39,65^, all of which have been reported in various models of demyelination^66–68^. While the relationship between these phenotypes remains unclear, recent studies suggest DAM are highly proliferative before becoming senescent^65^ and that inhibiting microglial proliferation limits DAM and senescent microglia accumulation^69^. Indeed, we found that almost all commonly associated DAM genes were upregulated in lesions at 20 dpl compared to non-lesioned control white matter (**Extended Data Fig. 7b**). Together, it is conceivable that in response to myelin damage, microglia proliferate and enter the DAM/WAM activation state to process the debris. However, a subset of microglia may proliferate too vigorously, leading to the initiation of senescence programming. While SCs are successfully cleared from demyelinated lesions in young mice, the aged immune system may be less effective, leading to inefficient remyelination.

In conclusion, this study provides an important step to understand how demyelination promotes the accumulation of SCs in lesions, and how, especially with age, these cells limit remyelination through SASP production. Future studies are still needed to elucidate the exact contributions of individual lingering SASP factors in remyelination with age. Our study suggests eliminating SCs alone may not be sufficient to enhance remyelination with age, and that inhibiting lingering SASP activity, such as CCL11, may be a better strategy to improving the lesion environment for remyelination. This approach may have therapeutic potential for reversing the aging phenotype and enhancing the remyelination process in MS.

## METHODS

### Mice

All transgenic mice were maintained on a C57BL/6 background, and experiments were performed according to the protocol approved by the Institutional Animal Care and Use Committee at Georgetown University. Heterozygous p16-tdTomato mice (B6J.Cg-*Cdkn2a^tm4Nesh^*/Mmnc) were obtained through the NIH Mutant Mouse Resource & Research Center^26^. INK-ATTAC mice were obtained from Unity Biotechnology. C57BL/6 mice were purchased from Jackson Labs (strain 000664). Mice were kept in home-cages and maintained on a 12-hour-light/12-hour-dark cycle with food and water ad libitum until the comparable human age of young adult (2-3 months), middle aged (12 months), or aged adult (>18 months). Mice of both sexes were used for all experiments.

### Focal Demyelination

Focal demyelinating lesions were induced as previously described^27^. In brief, 1 ul of 1% L-α-Lysophosphatidylcholine (LPC, MilliporeSigma #L4129) in PBS was injected via pulled glass needle attached to a Hamilton syringe between the T11-T12 vertebrae into the ventral white matter columns of the mouse spinal cord. Remyelination was measured at 20 dpl, except where noted in 18-month mouse experiments at 30 dpl and 60 dpl.

### Treatments

#### AP20187

INK-ATTAC mice were treated twice a week intraperitoneally with AP20187 (2 mg/kg, MedchemExpress B/B homodimerizer, #HY-13992) or with vehicle (10% DMSO, 40% peg400, 5% tween 80, 45% saline) at equal volumes (10ul/g bodyweight) from day of lesion until final timepoint.

#### Dasatinib/Quercetin

12 month old C57BL/6 mice were treated three consecutive days twice, one week apart, via oral gavage with dasatinib monohydrate (5 mg/kg, ThermoFischer 462320010) and quercetin hydrate (50 mg/kg, ThermoFischer 174070100) or vehicle (10% etoh, 30% peg400, 60% phosphal 50 PG) at equal volumes (5ul/g bodyweight) starting on day 8 post lesion.

#### CCL11 Neutralizing Antibody

12 week old INK-ATTAC mice were treated twice a week intraperitoneally with CCL11 neutralizing antibody (50ug/kg, rndsystems #MAB420-100), control rat IgG2a isotype (50ug/kg, rndsystems #MAB006), or recombinant CCL11 with AP treatment (10ug/kg, rndsystems #420-ME-020/CF, and 2mg/kg, #HY-13992, respectfully) at equal volumes (10ul/g bodyweight) from day of lesion until final timepoint.

### Senescence Associated β-Galactosidase Staining

SA B-gal staining was performed following the manufacturer’s protocol (cell signaling #98605), with modifications. In brief, tissue was drop fixed in 4% paraformaldehyde in PBS for 2 hours prior to 30% sucrose and then processed and sectioned as with IHC. After thaw, sections were processed per manufacturers protocol before the addition of nuclear fast red (Bioenno #003034) for counterstaining and PBS rinse before coverslips.

### In Vivo Imaging System (IVIS)

At 5 dpl, p16-tdtom lesioned, p16-tdtom sham lesioned, and C57BL/6 wild-type control lesioned mice were anesthetized and bioluminescence was imaged using the IVIS Lumina III In Vivo Imaging System.

### Real Time qPCR

To identify and dissect out the lesion, 500ul 1% neutral red dye in PBS was injected intraperitoneally 2 hours prior to cardiac perfusion with ice cold 1X PBS^27^. RNA was isolated using the Direct-zol RNA Miniprep Kits (Zymo Research #R2050) and reverse transcribed to cDNA using the iScript gDNA Clear cDNA Synthesis Kit (Biorad #1725034). RT-qPCR was performed with SsoAdvanced Universal SYBR Green Supermix (Biorad #1725271). Primers used were purchased from Bio-Rad: *Rpl13a* (qMmuCED0040629)*, Cdkn2a* (qMmuCED0038108)*, Rb1* (qMmuCID0005286)*, Tert* (qMmuCID0018719)*, Apoe* (qMmuCED0044813)*, Trem2* (qMmuCID0020213)*, Cdkn1a* (qMmuCED0025027)*, E2f1* (qMmuCID0005113)*, Trp53* (qMmuCID0006264). Expression levels were expressed using the 2^−ΔΔCt^ method normalized to non-lesion tissue.

### Immunohistochemistry

Mice were cardiac perfused with ice cold 4% paraformaldehyde in PBS and dissected spinal cords were post fixed overnight at 4C, followed by 30% sucrose (w/v) in PBS overnight at 4C, before being embedded and frozen in Tissue-Tek O.C.T. compound (Sakura). Lesions were sectioned on a Leica CM1860 cryostat at 12 um. For staining, slides were thawed and dried for 1 hour at room temperature (RT) prior to being subsequently washed in TBST (1X TBS, 0.05% Tween), TBS, and permeabilization buffer (1X TBS, 1% Triton X-100). Slides were then blocked for 1 hour at RT in blocking/antibody buffer (5% Donkey serum, 1%BSA, 0.4% Triton X) before being stained overnight at 4C with the following primary antibodies: mouse anti-APC (CC1) (Abcam #ab16794, 1:100), goat anti-CCL11 (R&DSystems #AF-420-SP, 1:50), rat anti-Cd11b (Biorad #MCA74G, 1:100), CD68 (Biolegend #137020, 1:100), rat anti-Clec7a (InvivoGen #mabg-mdect, 1:100), mouse anti-GFAP (Millipore Sigma #G3893, 1:500), rabbit anti-Iba1 (Fujifilm Wako #019-19741, 1:400), mouse anti-iNos (BD Biosciences #610329, 1:100), rat anti-MBP (Biorad #MCA409S, 1:500), mouse anti-Nkx2.2 (DSHB #74.5A5-c, 1:100), rabbit anti-Olig2 (Millipore Sigma #AB9610, 1:300), mouse anti-p16 (Abcam #ab54210, 1:500), mouse anti-p21 (Santa Cruz #sc6246, 1:500), rabbit anti-RFP (Rockland Immunochemicals #600-401-379, 1:1000), rat anti-Sall1 (Invitrogen #14-9729-82). Slides were then washed subsequently in TBST and TBS before being incubated for 1 hour in the dark at RT with the following fluorescent conjugated secondary antibodies: Alexa Fluor 488 (Invitrogen, 1:1000), Alexa Fluor 594 (Invitrogen, 1:500) or Cy3 (Jackson ImmunoResearch, 1:500), Cy5 (Jackson ImmunoResearch, 1:500) and Hoechst 33342 (Invitrogen, 1:20,000). Following incubation, the slides were washed again in TBST and TBS before being rinsed with ddH_2_O, dried, and mounted with Fluoromount-G mounting medium (Southern Biotech # 0100-01).

For CC1, Olig2, Nxk2.2, Sall1, and CCL11 staining, antigen retrieval was performed prior to staining by incubating the slides in boiling Antigen Retrieval solution (Vector #H-3301-250) for 30 minutes. For MBP staining, slides were incubated in pre-cooled methanol for 10 minutes prior to blocking.

### Imaging and Quantification

Slides were imaged using a Zeiss LSM 800 confocal microscope. Samples were blinded by cage number and ear-tag. For each sample, 3 lesion sections and 1 non-lesion section were imaged. For analysis, lesions were identified using the DAPI channel and tracing the hyper-nucleated region of the white matter. Percent areas were calculated in FIJI (ImageJ) for each image as follows: background was subtracted, each channel was thresholded to pre-determined values based on the channel and experiment, and “% area” of region of interest (pre-traced lesions) was measured. Cells were counted in FIJI (ImageJ) using the “cell counter” plugin and then counts per mm^2^ were calculated. 3D reconstruction analysis was done using Imaris microscopy image analysis software (Oxford Instruments).

### Scanning Electron Microscopy and g-ratio

To identify and dissect out the lesion in a mouse spinal cord, 500ul 1% neutral red dye in PBS was injected intraperitoneally 2 hours prior to cardiac perfusion with ice cold fixation buffer (2.5% glutaraldehyde, 1% paraformaldehyde, in 0.2 M sodium cacodylate buffer (Electron Microscopy Sciences). Lesion tissues were then dissected out and cut into 2mmx2mm sections, washed with cold cacodylate buffer (2mM calcium chloride), and then incubated in osmium solution for 1 hour on ice. Tissues were then washed with ddH_2_O, incubated with 1% thiocarbohydrazide (TCH) solution (Ted Pella) for 20 minutes at RT, washed again with ddH_2_O, incubated with 2% osmium tetroxide for 30 minutes at RT, washed with ddH_2_O, and then incubated with 1% uranyl acetate at 4C overnight. Next, lesioned tissues were then washed with ddH_2_O and incubated in pre-heated lead aspartate solution at 60C for 30 minutes, followed by ddH_2_O wash, and dehydration with ice-cold freshly prepared 50%, 70%, 85%, 95%, 100% ethanol solutions, and propylene oxide. The tissues were then embedded in resin and incubated at 60C for 48 hours.

For image acquisition, ultrathin sections (120 nm) containing spinal cord lesions were mounted in silicon wafers and observed with a Teneo LV FEG scanning electron microscope (FEI, Thermo Fisher Scientific). For optimal results, we used the optiplan mode (high-resolution) equipped with an in-lens T1 detector (Segmented A+B, working distance of 8 mm). Low-magnification images (600X) of the entire spinal cord section were first taken to delineate the lesioned site, and then high magnification tile images with multiple captures of our regions of interest (10,000X) were taken to ensure comprehensive coverage of the demyelinated lesion area using 2 kV and 0.4 current landing voltage. For quantification, 2-3 mice (>300 remyelinated axons) from each group were measured for axon diameters and g-ratios (axon diameter/myelin diameter) using MyelTracer^28^.

### SASP Multiplex

To identify and dissect out the lesion, 500ul 1% neutral red dye in PBS was injected intraperitoneally 2 hours prior to cardiac perfusion with ice cold 1X PBS^27^. Spinal cords were extracted and lesion and distant contralateral non-lesion tissue were microdissected out and flash frozen on dry ice. Tissue was homogenized using RIPA buffer with 1% protease inhibitor (Millipore Sigma #P8340) and handheld homogenizer (Fisher Scientific #12-141-361). Following centrifugation, supernatant was moved to new tube and concentrations were determined via Bradford assay. 100ul from each sample was then normalized to contain the same amount of protein before being frozen and sent for bead-based multiplex analysis (Eve Technologies, 32-Plex Discovery Assay #MD32).

### Bulk RNA Sequencing

To identify and dissect out the lesion, 500ul 1% neutral red dye in PBS was injected intraperitoneally 2 hours prior to cardiac perfusion with ice cold 1X PBS^27^. Spinal cords were extracted and lesion and distant contralateral non-lesion tissue were microdissected out and flash frozen on dry ice. RNA was isolated using the Direct-zol RNA Miniprep Kits (Zymo Research #R2050) and sent to Azenta Life Sciences (South Plainfield, NJ, USA) for RNA quality check, library preparation, sequencing and analysis. RNA samples were quantified using Qubit 3.0 Fluorometer (Life Technologies, Carlsbad, CA, USA) and RNA integrity was checked using Agilent TapeStation 4200 (Agilent Technologies, Palo Alto, CA, USA). ERCC RNA Spike-In Mix kit (ThermoFisher #4456740) was added to normalized total RNA prior to library preparation following manufacturer’s protocol. RNA sequencing libraries were prepared using the NEBNext Ultra II RNA Library Prep for Illumina using manufacturer’s instructions (NEB, Ipswich, MA, USA). Briefly, mRNAs were initially enriched with Oligod(T) beads. Enriched mRNAs were fragmented for 15 minutes at 94 °C. First strand and second strand cDNA were subsequently synthesized. cDNA fragments were end repaired and adenylated at 3’ends, and universal adapters were ligated to cDNA fragments, followed by index addition and library enrichment by PCR with limited cycles. The sequencing libraries were validated on the Agilent TapeStation (Agilent Technologies, Palo Alto, CA, USA), and quantified by using Qubit 3.0 Fluorometer (Invitrogen, Carlsbad, CA) as well as by quantitative PCR (KAPA Biosystems, Wilmington, MA, USA).

The sequencing libraries were clustered on a lane of a NovaSeq 6000 S4 flowcell. After clustering, the flowcell was loaded on the Illumina instrument according to manufacturer’s instructions. The samples were sequenced using a 2×150bp Paired End (PE) configuration. Image analysis and base calling were conducted by the Control software. Raw sequence data (.bcl files) generated the sequencer were converted into fastq files and de-multiplexed using Illumina’s bcl2fastq 2.17 software. One mismatch was allowed for index sequence identification. After investigating the quality of the raw data, sequence reads were trimmed to remove adapter sequences. The trimmed reads were mapped to the mouse reference genome GRCm38 available on ENSEMBL using the STAR aligner v.2.5.2b. The STAR aligner is a splice aligner that detects splice junctions and incorporates them to help align the entire read sequences. BAM files were generated as a result of this step. Unique gene hit counts were calculated by using feature Counts from the Subread package v.1.5.2. Only unique reads that fell within exon regions were counted. After extraction of gene hit counts, the gene hit counts table was used for downstream differential expression analysis. Using DESeq2, a comparison of gene expression between the groups of samples was performed. The Wald test was used to generate p-values and Log2 fold changes. Genes with adjusted p-values < 0.05 and absolute log2 fold changes > 1 were called as differentially expressed genes for each comparison.

### Multiplexed Error-Robust Fluorescence In Situ Hybridization (MERFISH)

To identify and dissect out the lesion, 500ul 1% neutral red dye in PBS was injected intraperitoneally 2 hours prior to cardiac perfusion with ice cold RNA-ase free 1X PBS^27^. Spinal cords were flash frozen in isopentane and embedded in OCT on dry ice. For MERFISH processing, Vizgen’s fresh frozen tissue sample preparation protocol was followed. Briefly, spinal cord sections were sliced at 10 μm. Tissue slices were mounted on Vizgen’s bead-coated functionalized coverslip. Once adhered to the coverslip, the samples were fixed (4% PFA in 1X PBS, 15 min, room temperature) and washed (3x, 5 min, 1X PBS). Then samples were photobleached for 3 h in 70% EtOH (Vizgen’s photobleacher) followed by an overnight incubation to permeabilize the tissue. Following permeabilization samples were incubated for 30 min in Formamide Wash Buffer (30% deionized formamide (Sigma, S4117) in 2× SSC (Thermo Fisher, AM9765)) followed by the addition of the gene library mix for the hybridization step (48h, 37°C). Samples were then washed twice with Formamide Wash Buffer (2x, 30 min, 47°C), and then embedded in a gel embedding solution (0.5% of 10% w/v ammonium persulfate, 0.05% TEMED, 4% 19:1 acrylamide/bis-acrylamide solution, 0.3M NaCl, 0.06M Tris PH8), followed by overnight incubation with tissue clearing solution (2X SSC, 2% SDS, 0.5% v/v Triton X-100, and proteinase K 1:100) at 37°C. Finally, samples are washed, incubated with DAPI and polyT solution (15 min RT), and washed with formamide wash buffer (10 min RT).

The MERSCOPE 500 gene imaging kit was activated by adding 250 μl of Imaging Buffer Activator (Vizgen, #203000015) and 100 μl of RNAse Inhibitor (New England BioLabs, M0314L). 15 ml of mineral oil was overlaid on top of the imaging buffer through the activation port. After instrument priming and chamber assembly as per MERSCOPE user guide, a 10x low resolution mosaic was acquired for imaging area selection. Raw data was decoded using Vizgen’s analyzing pipeline incorporated in the MERSCOPE. Vizgen’s postprocessing tool (Vizgen, Cambridge, MA) was then applied to obtain the cell segmentation based on the DAPI staining by using the CellPose algorithm. Segmentation was performed on the middle Z plane (3rd out of 7) and cell borders were propagated to z-planes above and below. MERFISH processed data was analyzed in RStudio using Seurat 4.1.3, R 4.2.2 and custom-made scripts. Cell filtering was applied to the dataset to remove cells <50 µm3 and <10 unique transcripts and <10 transcript counts. Cell gene expression per cell was then normalized to each cell’s volume and the mean RNA per sample. To perform cell clustering we followed a modified Seurat pipeline. We performed principal component analysis (PCA) with 400 genes as the variable features followed by dimensionality analysis with a resolution of 1.45 and 20 dimensions. Dimensionality reduction was performed with Uniform Manifold Approximation and Projection (UMAP). Cell clusters were manually annotated according to the expression of widely used cell-type specific gene markers as well as spatial distribution.

### Re-analysis of snRNA-sequencing data

The published single nucleus dataset of the human MS brain (PRJNA544731) was used to validate the expression of the *p16/Cdkn2a* gene. The ambient RNA, a pervasive artifact in single nucleus datasets, was removed from the raw data using CellBender (v0.3.0, https://github.com/broadinstitute/CellBender)^29^, specifically the cellbender-remove background function including 50,000 droplets (total-droplets-included 50,000). The subsequent quality control, dimensionality reduction and clustering were performed using Seurat (v5.0, https://github.com/satijalab/seurat)^30^, in the R statistical environment (v4.1.2). The filtered CellBender output was read into Seurat using the Read10X_h5() function and a SeuratObject was created using the CreateSeuratObject() function, retaining only the cells expressing at least 200 genes and genes expressed in at least 3 cells. The remaining 26,201 genes and 49,995 cells were further filtered to exclude doublets/multiplets by removing cells with high gene counts (>7500) and dead cells by removing cells with a high proportion of mitochondrial genes (>10%). The dataset was normalized using the SCTransform() function according to the binomial regression model^31^ (Highly variable features = 3000, regressed nCount and mitochondrial genes). The dimensionality reduction was performed using RunPCA(), FindNeighbors() (Dimensions = 20) and FindClusters() functions. Twenty-five PCs were used for downstream analyses as determined by the PCA elbow plot (min.dist = 0.3). The optimal clustering resolution was determined to be 0.5 using the Clustree package (v0.5.1, https://github.com/lazappi/clustree)^32^. The FindClusters() function was used to cluster the dataset at 11 resolutions ranging from 0 to 1, separated by 0.1 All resolutions were plotted on a tree generated through by clustree() function and 0.5 was identified as the most stable clustering level. The microglia (n=1815) were subsetted using the subset() function based on *TMEM119*/*P2RY12* expression and clustered again as described above. The differential gene expression was determined using the FindMarkers() function. The SeuratObject was converted to an h5ad object using SeuratDisk by converting the RDS object to an h5Seurat object using the Saveh5Seurat() function and converting the h5Seurat object to an h5ad object using the convert() function (Destination = h5ad). The h5ad file was read into a Jupyter notebook (v6.0.3) running a python environment (Python 3.8.3) using Scanpy (v1.6.0, https://github.com/scverse/scanpy)^33^. The final clustering was projected onto a UMAP using the pl.umap() function and the genes were plotted using the pl.violin() function.

### Human Samples

Samples were collected at the Rocky Mountain Multiple Sclerosis Center Biorepository at University of Colorado Anschutz Medical Campus as follows. Blood is collected as part of standard of care procedures in two 15 mL glass vacutainers containing 1.5 mL 3.8% Sodium Citrate solution. Both vacutainers are then pooled together in a 50 mL Leucosep tube (Greiner Bio-One) filled with 15 mL Lymphoprep (Stemcell). The Leucosep tube is then centrifuged at 1800 x g for 15 minutes at half brake. After centrifugation, plasma is then removed from above the mononuclear cell layer and transferred into a 15 mL conical. Plasma is then centrifuged at 500 x g for 10 minutes at full brake. Plasma is then aliquoted and transferred to −80 C for storage.

### ELISAs

CCL11 was measured in blood plasma from healthy controls and MS patients according to the manufacturer’s protocol (R&D Systems, #DTX00).

### Data Analyses

All statistics were performed in GraphPad Prism 10. Data is presented as mean ± SEM. Each value represents an individual biological sample, where technical replicates have been averaged. Statistical significance is reported as: ns (p>0.05), * (p<0.05), ** (p<0.01), *** (p<0.001), **** (p<0.0001).

## ACKNOWLEDGEMENTS

P.S.G. was supported by the Center for Neural Injury and Recovery (CNIR) program (NINDS/NIH T32NS041218). J.K.H. was supported by NIA/NIH (R21AG072327), NINDS/NIH (R01NS107523), and NMSS Harry Weaver Neuroscience Scholar Award (JF-1806-31381). S.S. was supported by the Rocky Mountain Multiple Sclerosis Center. We would like to thank Olga Rodriguez, MD, PhD from the Preclinical Imaging Research Laboratory (PIRL) at Georgetown University Medical Center for training on the In Vivo Imaging System (IVIS) imager. We would also like to acknowledge Eve Technologies and Azenta Life Sciences for their services. All illustrations were put together with BioRender.com.

## AUTHOR CONTRIBUTIONS

J.K.H. and P.S.G designed the study. P.S.G. performed all animal and human sample experiments and interpreted the data. Z.M. and S.H.L. contributed intellectually and assisted in animal experiments. V.D.L. processed the MERFISH data and developed the computational framework and performed associated analyses. P.S.G. and V.D.L. interpreted the MERFISH data. D.P.S. oversaw the MERFISH experiments. N.S. performed SEM imaging. S.Z. performed re-analysis of MS snRNAseq tissues and J.R.P. oversaw the snRNAseq re-analysis. S.S. and E.A. isolated and provided human blood plasma samples. P.S.G. wrote the original draft of the manuscript and J.K.H. co-edited the manuscript. All authors discussed the results and reviewed the manuscript. J.K.H. contributed to data interpretation and oversaw the study.

## DATA AVAILABILITY

All data are available in the main text or the supplementary materials. Any additional data that support the findings of this study are available from the corresponding author upon request.

---

**Extended Data Fig. 1.**
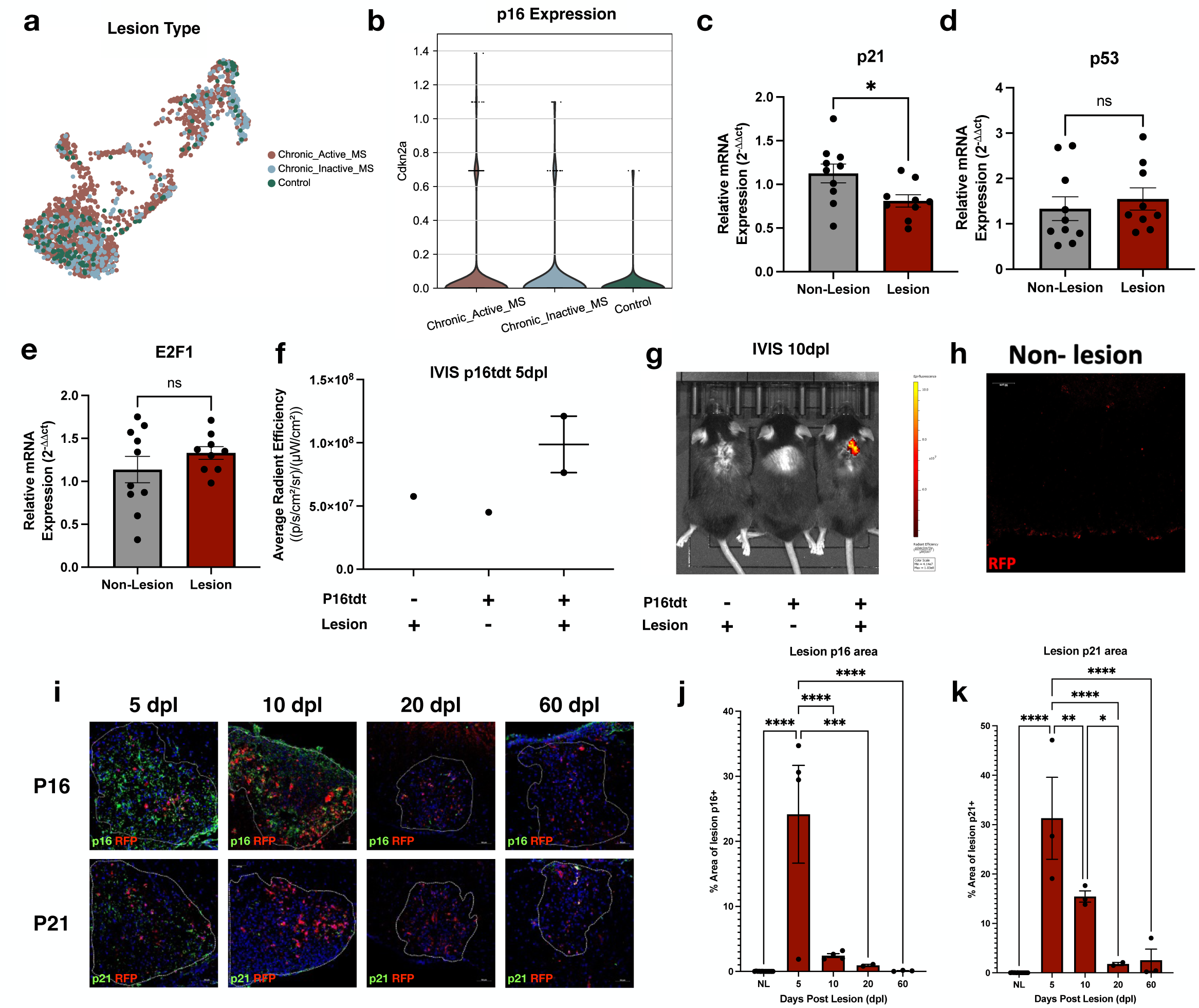
Senescent cells are observed within lesions following demyelination. **a,** Uniform manifold approximation and projection (UMAP) visualization of re-analyzed snRNA-sequencing from chronic active lesions, inactive lesions, and control samples (1815 total microglia). **b,** Violin plots describing expression levels of *p16/Cdkn2a* chronic active lesions, inactive lesions, and control samples (n=1321, 349, 145 microglia). **c-e,** RT-qPCR for selected senescence genes from dissected lesion (n= 9-10). **f,** Quantification of In Vivo Imaging System (IVIS) bioluminescence of controls and p16tdt lesions (n= 1-2 per group) at 5 dpl. **g,** IVIS bioluminescence of controls and p16tdt lesions at 10 dpl. **h,** Immunofluorescent staining of p16tdt+ cells in non-lesion white matter. **i,** Immunofluorescent staining of *p16* and *p21* throughout remyelination. **j,k** Quantification of g (n= 2-4 per timepoint). Data are mean ± SEM (**c-f,j,k**). RT-qPCR gene expression levels were expressed using the 2^−1ΔΔCt^ method normalized to non-lesion tissue (**c-e**). *P* values derived from two-tailed unpaired Student’s t-tests (**c-e**) or one-way ANOVA with Tukey corrected multiple comparisons (**j,k**).

**Extended Data Fig. 2.**
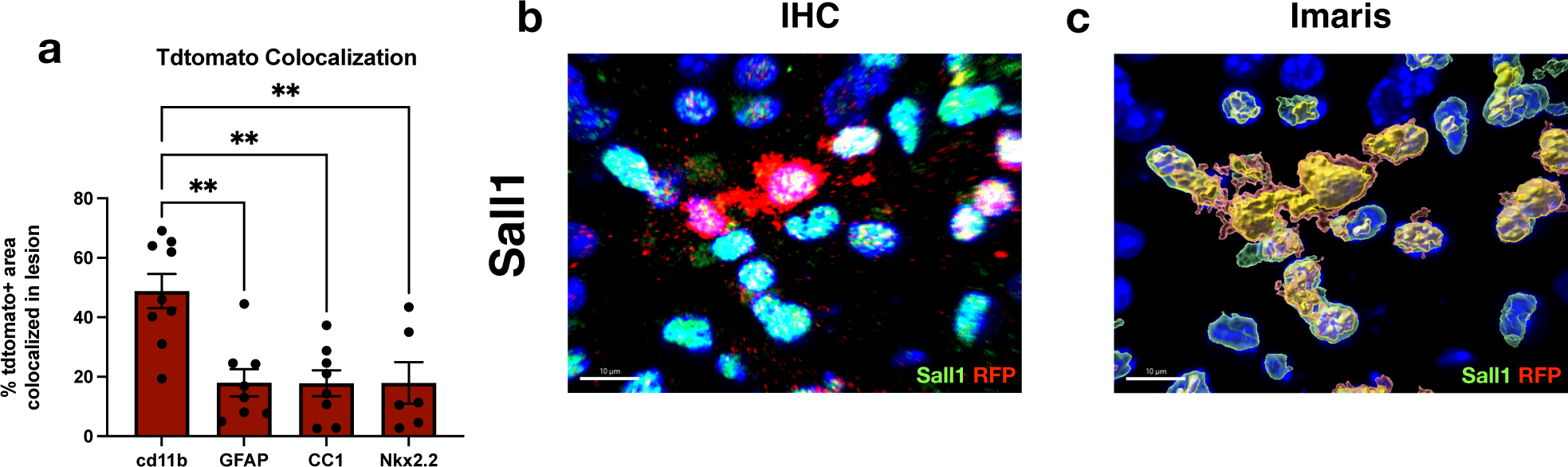
Confirming microglia colocalization. **a,** Quantification of colocalization of p16tdt and cell markers via ImageJ percent area (n= 6-8). **b-c,** Immunofluorescent staining and Imaris 3D reconstruction of p16tdt and *SALL1*. **d,** Heatmap depicting Damage associate microglia (DAM) gene expression in young lesions relative to young non-lesions (n=3). Color represents LogFC relative to young non-lesion and * indicates significantly differentially expressed. Data are mean ± SEM (**a**). *P* values derived from one-way ANOVA with Tukey corrected multiple comparisons (**a**) or Wald test (**d**).

**Extended Data Fig. 3.**
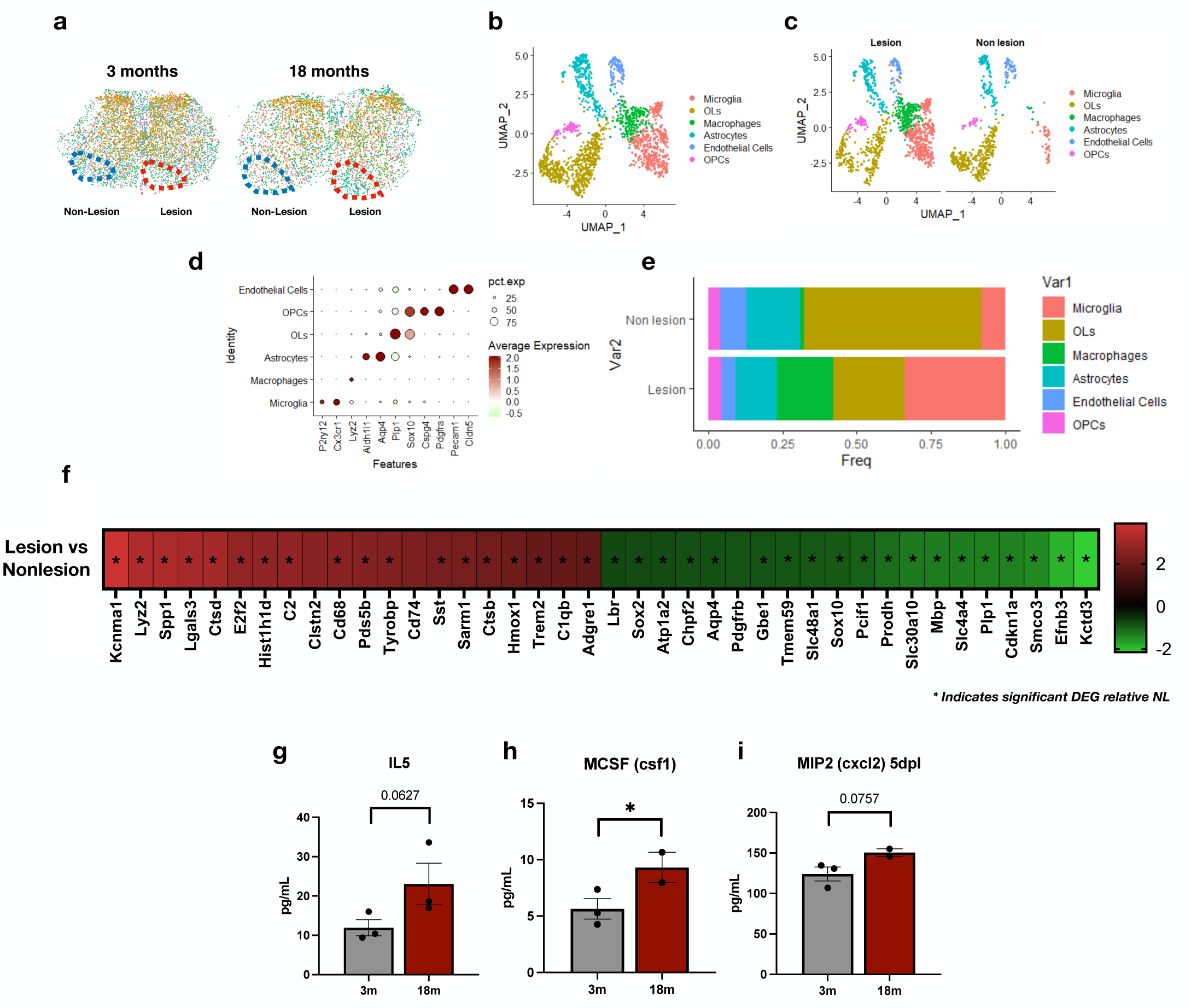
MERFISH analysis of 5 dpl lesion vs non-lesion and SASP factors. **a,** Demyelinated lesions and non-lesions were traced in young and aged mice at 5 dpl. **b,** Uniform manifold approximation and projection (UMAP) visualization of pooled lesions and non-lesions regardless of age. **c,** UMAP visualization of cell types in lesions and non-lesions (n=2 per group). **d,** Cell-type associated gene markers used to identify major cell types found in the spinal cord. **e,** Relative percentage of cells identified in lesions between lesions and non-lesions. **f,** Heatmap depicting top differentially expressed genes between lesions and non-lesions at 5 dpl. Color represents LogFC relative to non-lesion and * indicates significantly differentially expressed. **g-i,** Levels of selected SASP factors from Fig. 2g. Data are mean ± SEM (**g-i**). *P* values derived from two-way ANOVA with FDR corrected multiple comparisons (**g-i**) or the Wald test (**f**).

**Extended Data Fig. 4.**
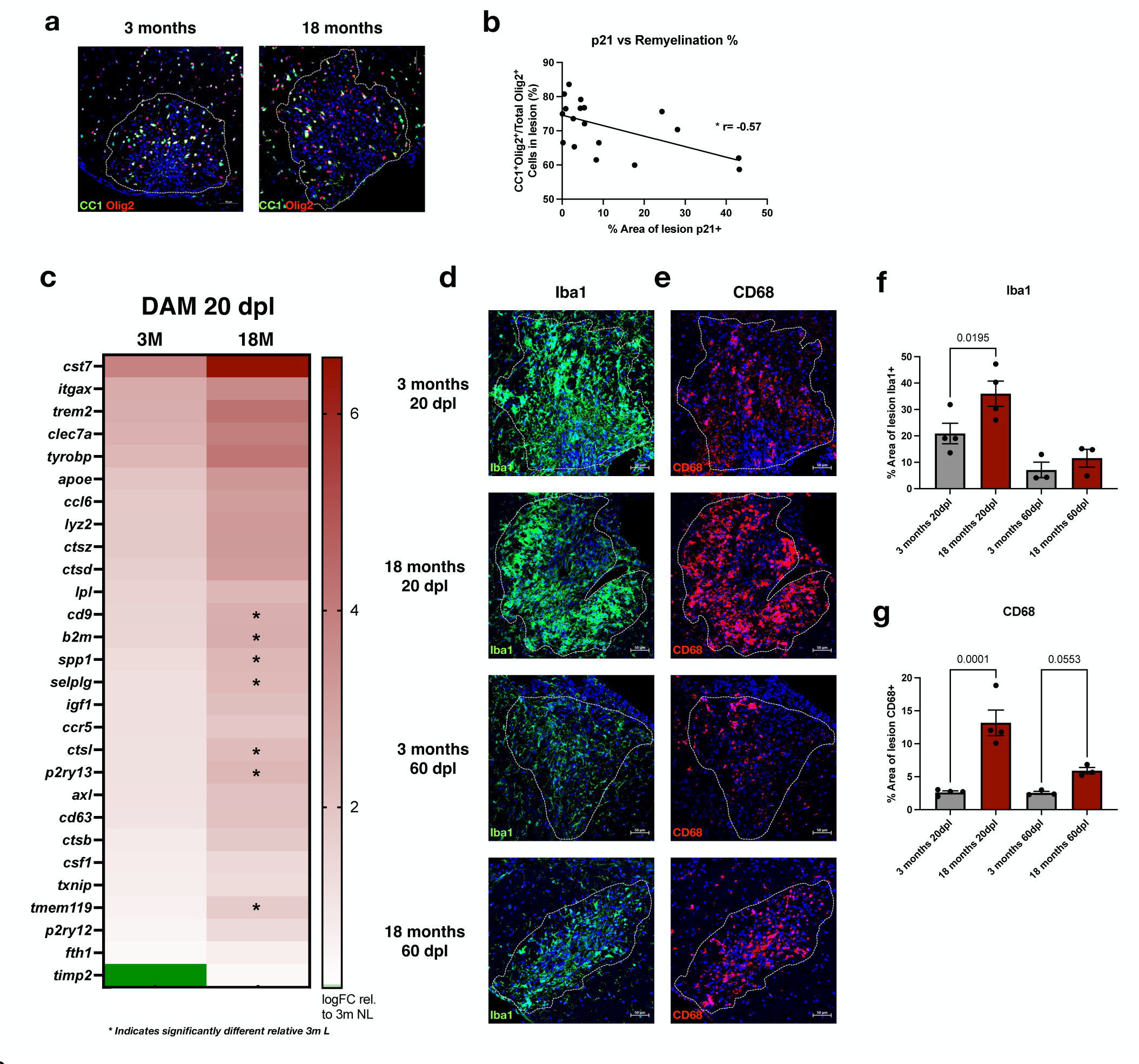
Neuroinflammation in young and aged lesions following demyelination. **a,** Immunofluorescent staining of mature oligodendrocytes and oligodendrocyte lineage cells with *CC1* and *Olig2* in young and aged lesions at 20 dpl. **b,** Correlation between p21+ staining and OPC differentiation efficiency in the lesions. **c,** Heatmap depicting Damage associate microglia (DAM) gene expression in young and aged lesions relative to young non-lesions (n=3). Color represents LogFC relative to young non-lesion and * indicates significantly differentially expressed from young lesions. **d,e,** Immunofluorescent staining of *Iba1* and *CD68* in young and aged lesions at 20 and 60 dpl. **f,g,** Quantification of *Iba1+* and *CD68+* cells in c,d (n= 3-4 per group). Data are mean ± SEM (**f,g**). *P* values derived from simple-linear regression (**b**), one-way ANOVA with Tukey corrected multiple comparisons (**f,g**) or Wald test (**c**).

**Extended Data Fig. 5.**
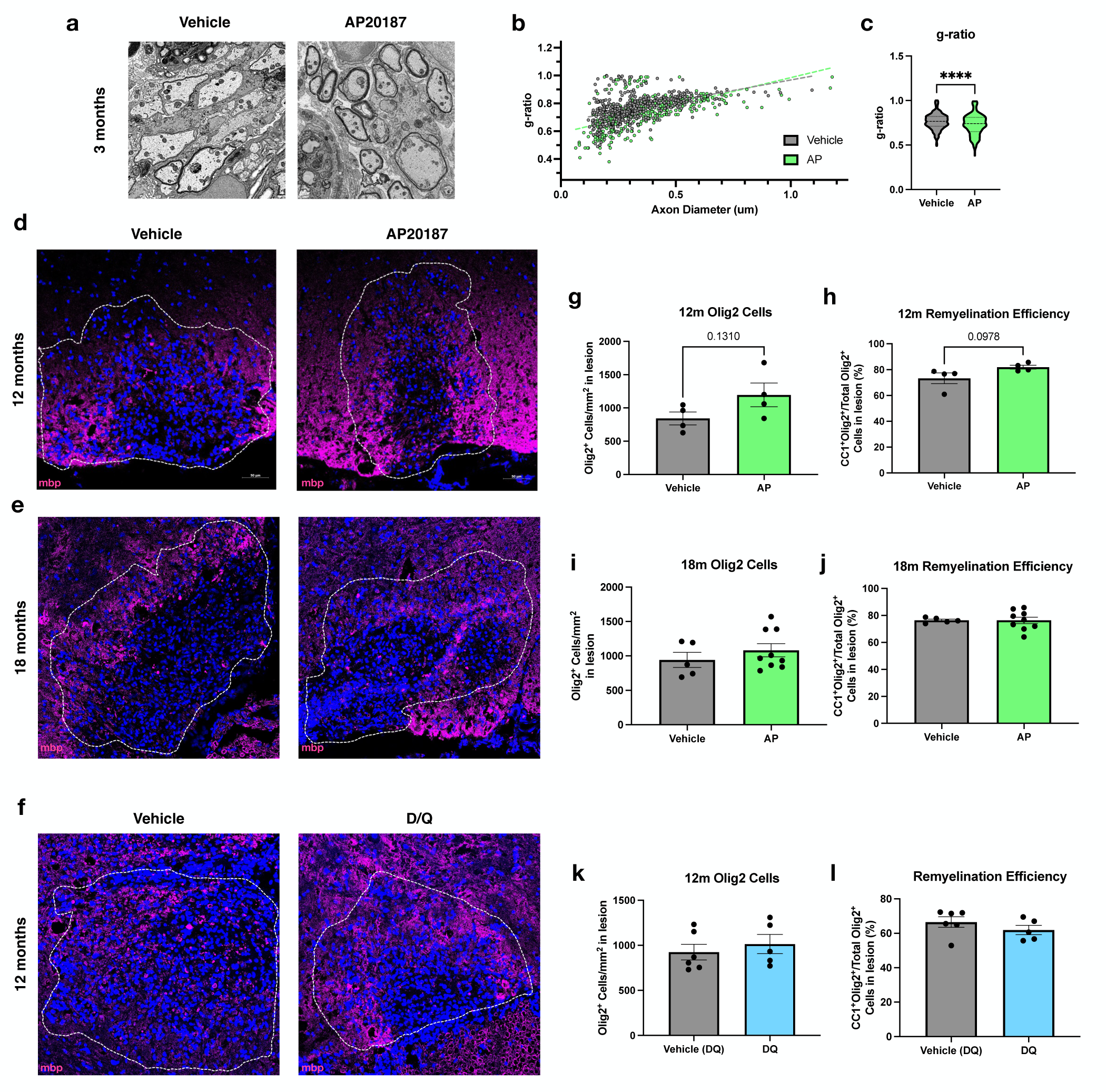
Additional markers of remyelination following AP and D/Q treatment. **a,** Scanning electron microscopy (SEM) images of remyelinated axons in young vehicle and AP treated lesions at 20 dpl (n= 2-3 mice per group). **b,c,** g-ratio’s of remyelinated axons counted from a (n= 300-400 axons per group). **d,** Immunofluorescent staining of myelin with *MBP* in vehicle and AP treated lesions at 20 dpl in middle aged (12 months) mice. **e,** Immunofluorescent staining of myelin with *MBP* in vehicle and AP treated lesions at 30 dpl in aged (18 months) mice. **f,** Immunofluorescent staining of myelin with *MBP* in vehicle and D/Q treated lesions at 20 dpl in middle aged (12 months) mice. **g-l,** Quantification of oligodendrocyte lineage cells (*Olig2+)* and OPC differentiation percentage of cells in Fig. 5g,j,m (n=4-9 per group). Data are mean ± SEM (**c,g-l**). *P* values derived from two-tailed unpaired Student’s t-tests (**c,g-l**).

**Extended Data Fig. 6.**
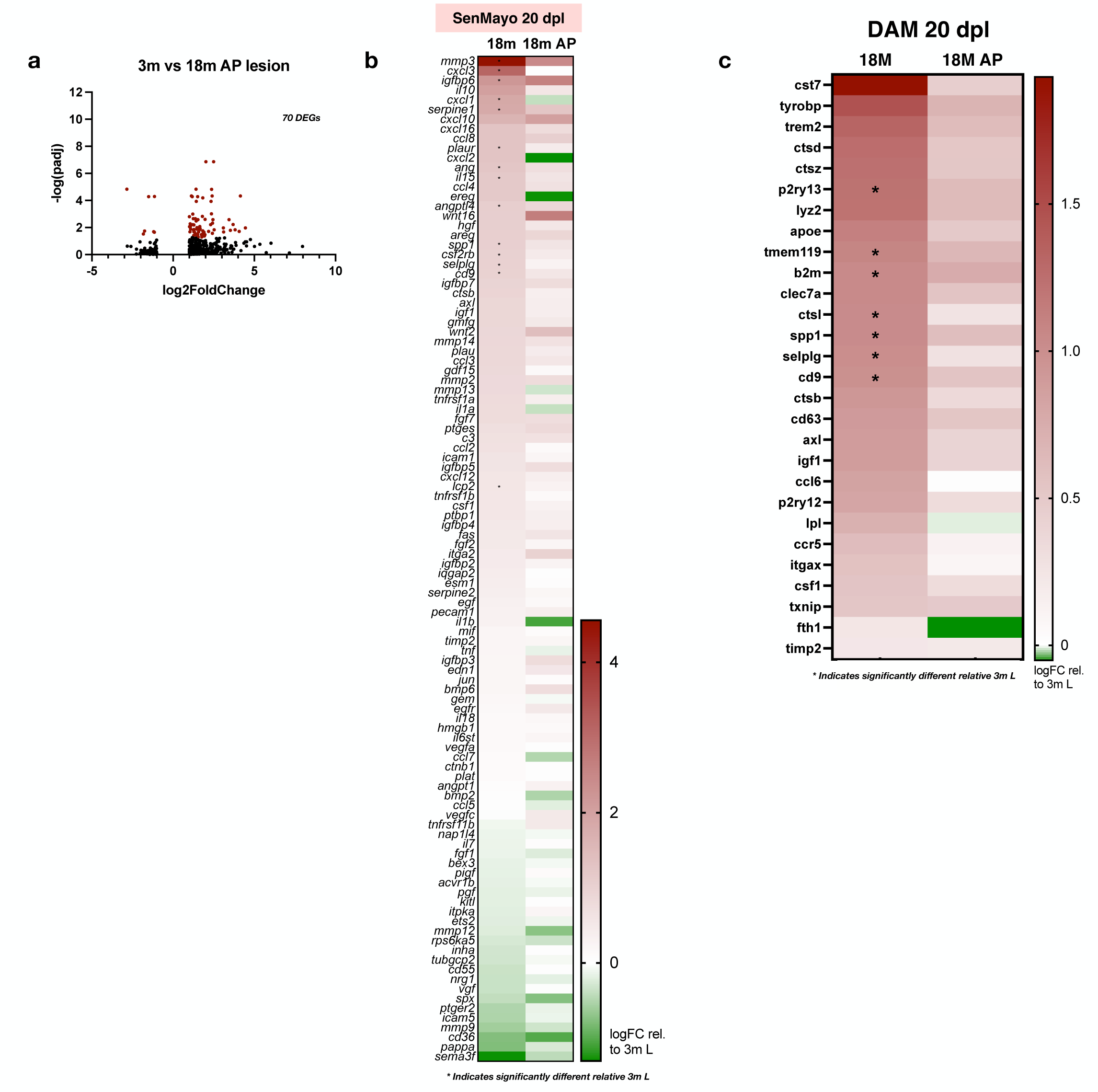
AP20187 treatment rejuvenates gene expression in aged demyelinated lesions. **a,** Volcano plot of significantly differentially expressed genes between young demyelinated lesions and aged lesions treated AP (n=3 per group) at 20 dpl. Genes with adjusted p-values <0.05 and absolute log2 fold changes >1 were called as differentially expressed genes for each comparison. **b,** Heatmap depicting SenMayo gene expression in aged vehicle and AP treated lesions relative to young lesions (n=3 per group). Color represents LogFC relative to young lesion and * indicates significantly differentially expressed **c,** Heatmap depicting Damage associate microglia (DAM) gene expression in vehicle treated aged lesions and AP treated aged lesions relative to young lesions (n=3). Color represents LogFC relative to young lesion and * indicates significantly differentially expressed from young lesions. *P* values derived from Wald test (**a,b,c**).

**Extended Data Fig. 7.**
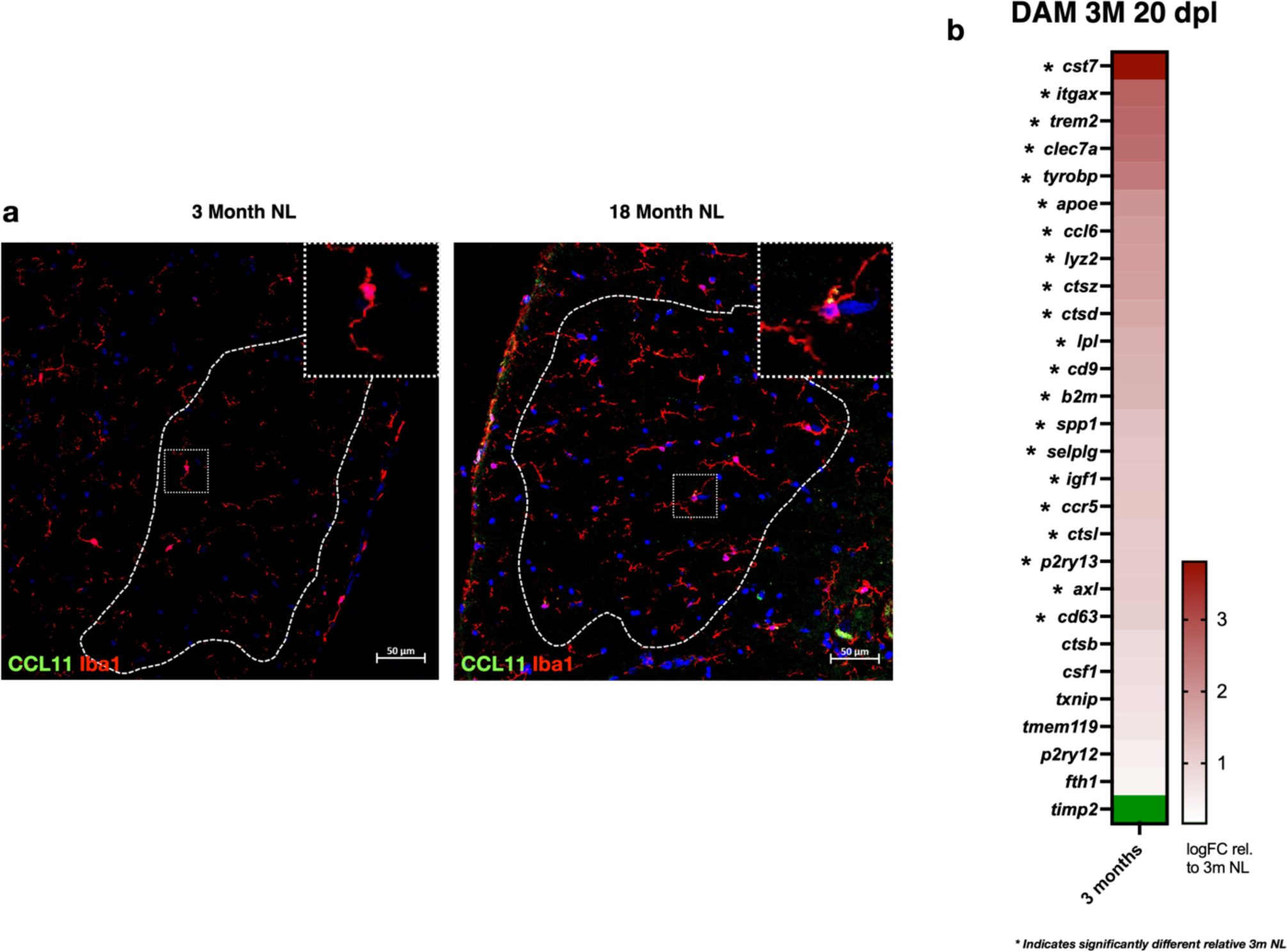
CCL11 expression is negligible in non-lesion white matter and proposed model of senescent cell genesis. **a,** Immunofluorescent staining of microglia (*Iba1*) and *CCL11* in young (3 months) and aged (18 months) non-lesion white matter. **b,** Heatmap depicting Damage associate microglia (DAM) gene expression in young lesions relative to young non-lesions (n=3). Color represents LogFC relative to young non-lesion and * indicates significantly differentially expressed. *P* values derived from Wald test (**b**).

